# Sedimentary ancient DNA reveals rhizosphere-like plant–microbe association signals in a 2-million-year-old Arctic ecosystem

**DOI:** 10.64898/2026.07.05.735348

**Authors:** Maria Landolfi, Nikolay Oskolkov, Edoardo Pasolli, Raphael Tiziani, Federica Villa, Tanja Mimmo, Eran Elhaik, Luigimaria Borruso

## Abstract

Plant–microbe interactions in the rhizosphere are central to nutrient cycling and ecosystem functioning. Sedimentary ancient DNA (sedaDNA) is a promising yet underexplored tool for reconstructing past microbial communities and investigating ecological interactions among plants, animals, and microorganisms. Here, we reanalyse the previously published Kap København Formation (Northern Greenland) sedaDNA dataset to move beyond taxonomic ecosystem reconstruction and test whether ancient sediments preserve structured, rhizosphere-compatible plant–microbe association signals. Our results show that this ancient boreal ecosystem hosted several rhizosphere-associated taxa, comparable to those in modern boreal soils. Several bacterial genera co-occurred repeatedly with specific plant families, forming a rhizosphere-like taxonomic core with predicted plant-growth-promoting traits related to nutrient acquisition, colonisation, and stress tolerance. Although sedaDNA co-occurrence cannot demonstrate direct symbiosis, the consistency of taxonomic, network, and functional signals suggests that ancient sediments preserve interconnected ecological structure. Our findings extend sedaDNA-based ecosystem reconstruction beyond taxonomy and provide a possibility for investigating plant–microbe association signals in deep time.

## Introduction

The rhizosphere, the narrow zone of soil influenced by root activity, is a hotspot of plant– microbe interactions that are fundamental to ecosystem functioning (Hinsinger et al., 2006). Rhizosphere microorganisms drive key biogeochemical cycles, including carbon (C), nitrogen (N), phosphorus (P), and sulphur (S), thereby influencing plant productivity, diversity, health, and community composition (Von Hippel et al., 2025; Zilber-Rosenberg & Rosenberg, 2008). This belowground interface is among the most biologically active environments on Earth, and its ecological importance is rooted in deep evolutionary history. Fossil evidence supports this view: for example, the Rhynie chert deposits in Scotland reveal plant–fungal symbioses resembling mycorrhizal structures dating back to the Devonian period (~407 Ma), suggesting that microorganisms may have facilitated the colonisation of land by early plants (Philippot et al., 2013). However, fossils alone provide limited information about the taxonomy and potential of ancient rhizosphere microorganisms, highlighting the need for complementary approaches that can directly access ancient genetic material.

In this context, ancient DNA (aDNA) has emerged as a powerful tool, transforming our ability to investigate the past beyond what can be inferred from modern DNA (Morozova et al., 2016). For instance, the recovery of aDNA from sediments (sedaDNA), though methodologically challenging, offers a powerful window into entire past ecosystems (Aldeias & Stahlschmidt, 2024; Fracasso et al., 2024; Jia et al., 2022; Willerslev & Cooper, 2005). Because SedaDNA contains genetic traces of all the organisms that once inhabited a given environment, it is an archive of ecological information that, theoretically, allows the reconstruction of entire ancient communities, despite potential challenges associated with ancient DNA, which increases for deep ancient collections. If these challenges are addressed, it is possible not only to identify species composition but also to investigate potential ecological association signals even when macrofossils and remains are absent (Ekram et al., 2025; Özdoğan et al., 2024; Willerslev & Cooper, 2005).

Within this broader framework, a particularly promising yet still largely unexplored application is the inference of ancient plant–microbe interaction networks, including rhizosphere microbiomes. By extending the focus beyond plant aDNA to include microbial aDNA, it becomes possible to explore how plant-associated communities were structured in deep time and to infer their functional traits relevant to growth promotion and biogeochemical cycling.

To demonstrate the potential of this approach, we used sedaDNA from the Kap København Formation (northern Greenland) to investigate potential plant-microbe association signals. This formation preserves genetic evidence of an approximately 2-million-year-old ecosystem that resembles those of a modern boreal forest (Kjær et al. 2022; Fernandez-Guerra et al. 2023). Previous works that reconstructed the biotic composition of this ecosystem, did not address whether ancient plant and microbial taxa were linked by recurring ecological associations. Moreover, most studies employing sedaDNA have focused mainly on the characterisation of individual plant, animal, or microbial components of ecosystems, without examining whether structured association signals among them can be recovered from ancient sedimentary archives.

Here, we reanalyse the previously published Kap København sedaDNA dataset and test whether ancient sediments preserve structured, rhizosphere-compatible association signals between plants and microorganisms.

In particular, we aimed to infer rhizosphere-like plant–microbe association signals in this ancient boreal-like ecosystem by studying co-occurrence patterns between plants and soil- and rhizosphere-associated bacteria, as well as their plant growth-promoting potential. We hypothesised that authenticated ancient microbial taxa with known soil or rhizosphere affinities would exhibit non-random co-occurrence with specific plant families (H1) and that microbial taxa involved in these association patterns would show a set of predicted plant-growth-promoting traits linked to plant growth promotion (H2).

## Results and discussion

The Kap København sediments preserve traces of ancient biodiversity (Fernandez-Guerra et al., 2023; Kjær et al., 2022). After filtering for low-quality reads, we retained an average of 333,935 reads per sample. Subsequently, we screened them to isolate the ancient microbial component through conservative authentication analyses of deamination signals and coverage evenness (see Materials and Methods). After this filtering, the final dataset comprised 180 ancient microorganisms and their relative abundances (Supplementary information).

Consistent with previously demonstrated terrestrial input (Fernandez-Guerra et al., 2023), we observed a high abundance of soil- and rhizosphere-associated taxa, particularly members of the order Rhizobiales. Specifically, the most abundant genera within Rhizobiales were *Rhizobium, Bradyrhizobium*, and *Sinorhizobium*, which include taxa with well-known N-fixing activity (Alzate Zuluaga et al., 2024; Black et al., 2012; Lindström & Mousavi, 2020; Saharan & Nehra, 2011). In addition, these genera are key components of the rhizosphere across diverse ecosystems, including arctic and boreal soils (Chu et al., 2010; Duan et al., 2022; Malard et al., 2019), with modern representatives known to form plant associations, enhance nitrogen availability, and act as plant growth-promoting rhizobacteria (PGPR) (Jaiswal et al., 2021; Saharan & Nehra, 2011; Vejan et al., 2016). Given the high abundance of plant- and soil-associated taxa, we further investigated potential relationships between bacteria and plants within the same samples. To assess whether plant–microbe associations were structured beyond random co-occurrence, we quantified plant–microbe co-occurrences across samples. This allowed us to test whether plant taxa systematically co-occurred with specific microbial groups while acknowledging that depositional dynamics may also contribute to co-distribution patterns. This analysis identified 317 statistically significant plant–microbe co-occurrences (FDR-adjusted p-value < 0.05; Fig. 2), primarily involving soil- and plant-associated taxa, including *Nocardia, Streptomyces, Rhodospirillum, Mycobacterium*, and several genera within the order Rhizobiales. To explore how these co-occurrences were distributed across different stratigraphic units, we next examined the modular structure of each bipartite network. Analysis of network modules for total co-occurrences (Fig. 3–5) revealed that the highest number of significant co-occurrence links was observed in Unit B3 samples. Unit B2 exhibited a more generalist pattern (Fig. 4), with co-occurrences more evenly distributed across the network, similar to Unit B3 (Fig. 5). Most of the associations were concentrated in the module located at the bottom right, which included genera commonly associated with plants, such as *Cereibacter, Ensifer, Streptomyces*, and *Sinorhizobium*. These microbes primarily co-occurred with members of the Boraginaceae, Brassicaceae, and Saxifragaceae families. By contrast, other genera, including *Mycobacterium* and *Mycolicibacterium*, were associated with plants from the Celastraceae, Cystopteridaceae, Papaveraceae, and Rosaceae families. While direct interactions between these genera and plant taxa have not been documented previously, we identify a novel co-occurrence pattern between them, which is plausible given the widespread presence of *Mycobacterium* and *Mycolicibacterium* in soil and their occupation of diverse ecological niches (Pereira et al., 2020; Walsh et al., 2019).

**Figure 1.**
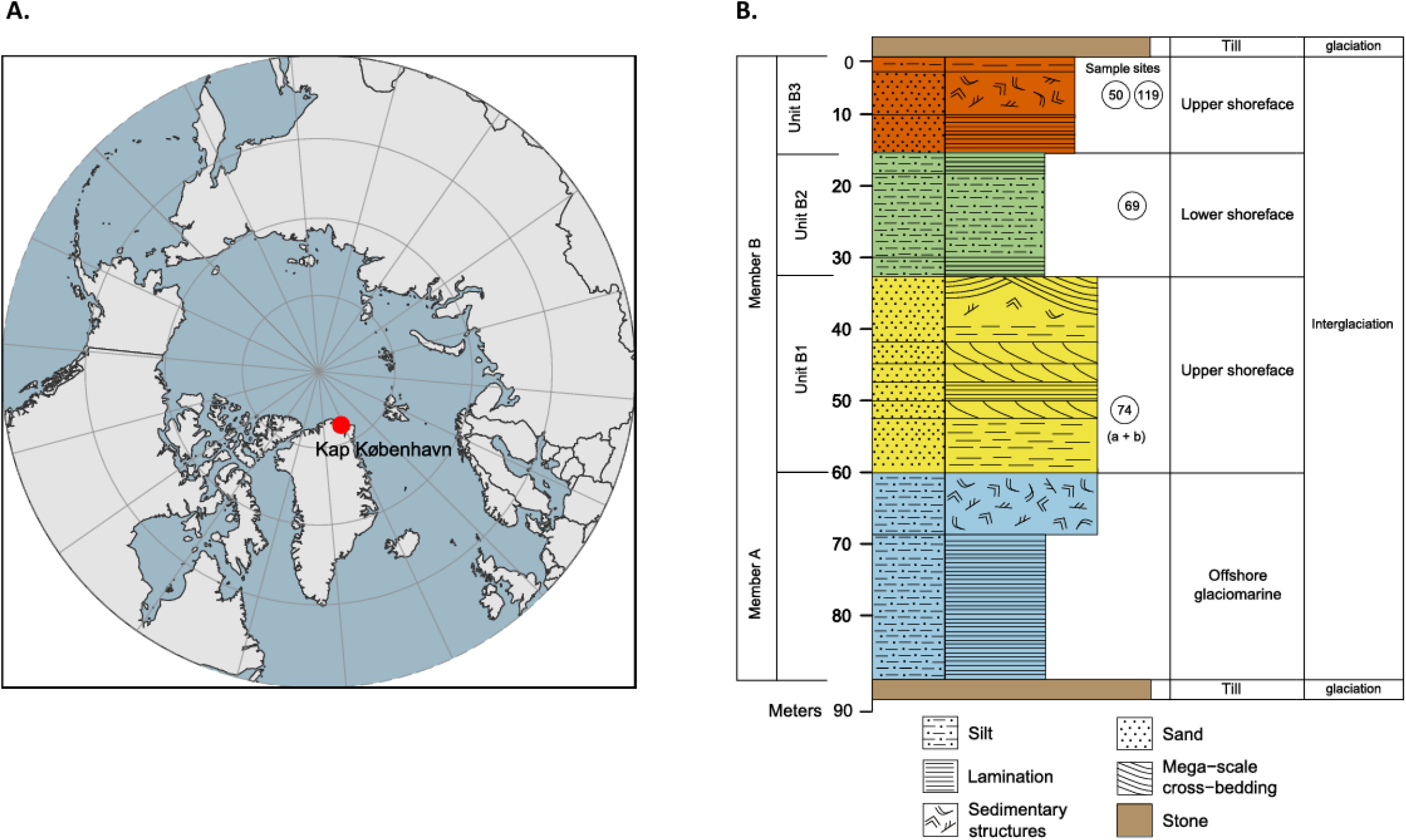
Geographical location and depositional sequence. A. Location of Kap København Formation in North Greenland at the entrance to the Independence Fjord (82° 24⍰ N 22° 12⍰ W). B. Glacial–interglacial division of the depositional succession of clay Member A and units B1, B2 and B3 constituting sandy Member B. Circled numbers indicate the sampling sites. Modified from Kjær et al. (2022).

**Figure 2.**
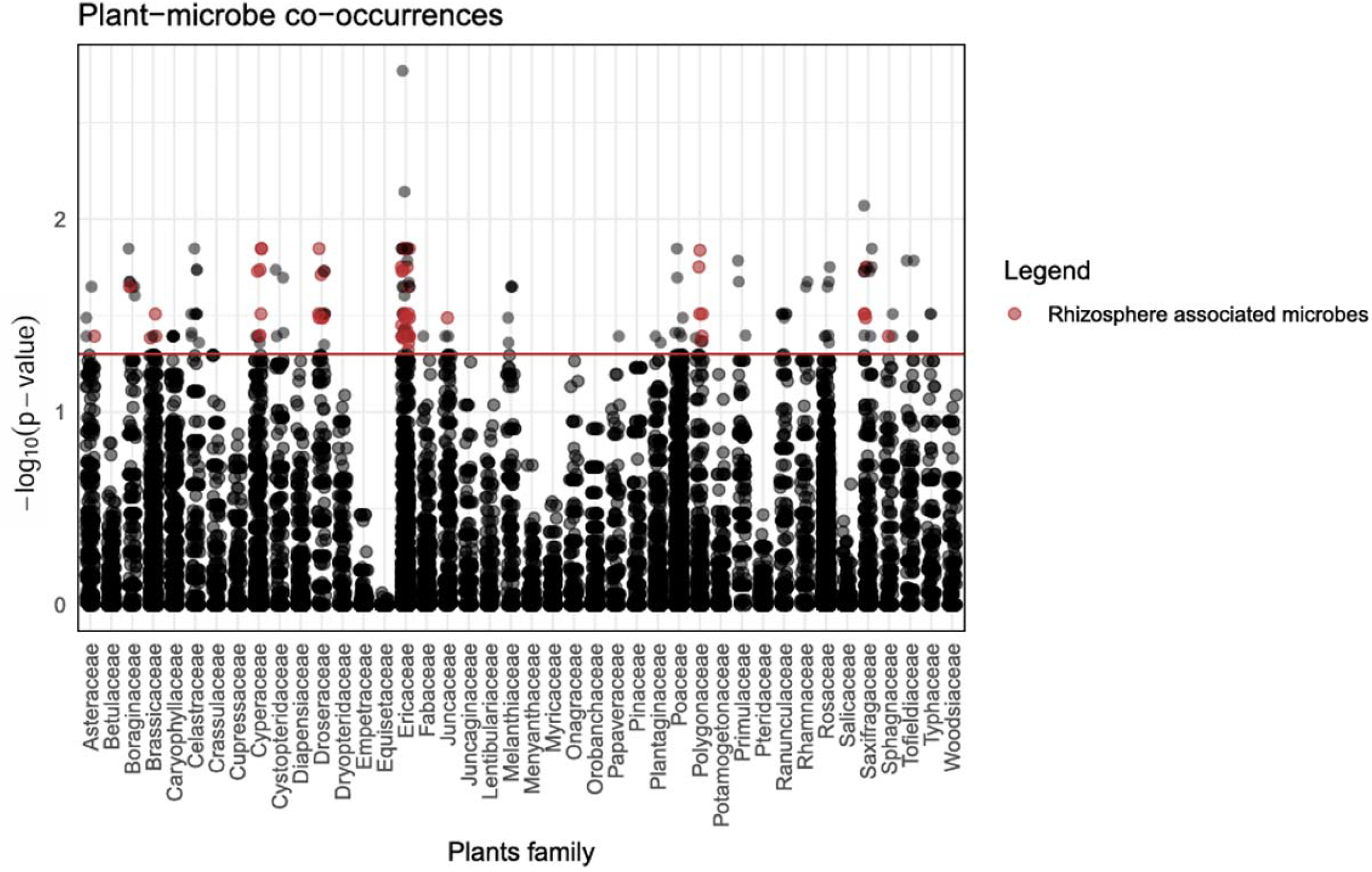
Vertical dotplot showing the total plant-microbes co-occurrences. Each point represents an association between a plant family (x-axis) and a microbial taxon, with the strength of the association indicated by –log_10_(p-value) on the y-axis. The red horizontal line marks the significance threshold (FDR-adjusted p-value < 0.05). Red points indicate association involving nitrogen-fixing bacteria, gray points indicate other significant associations, and black points represent non-significant associations.

**Figure 3.**
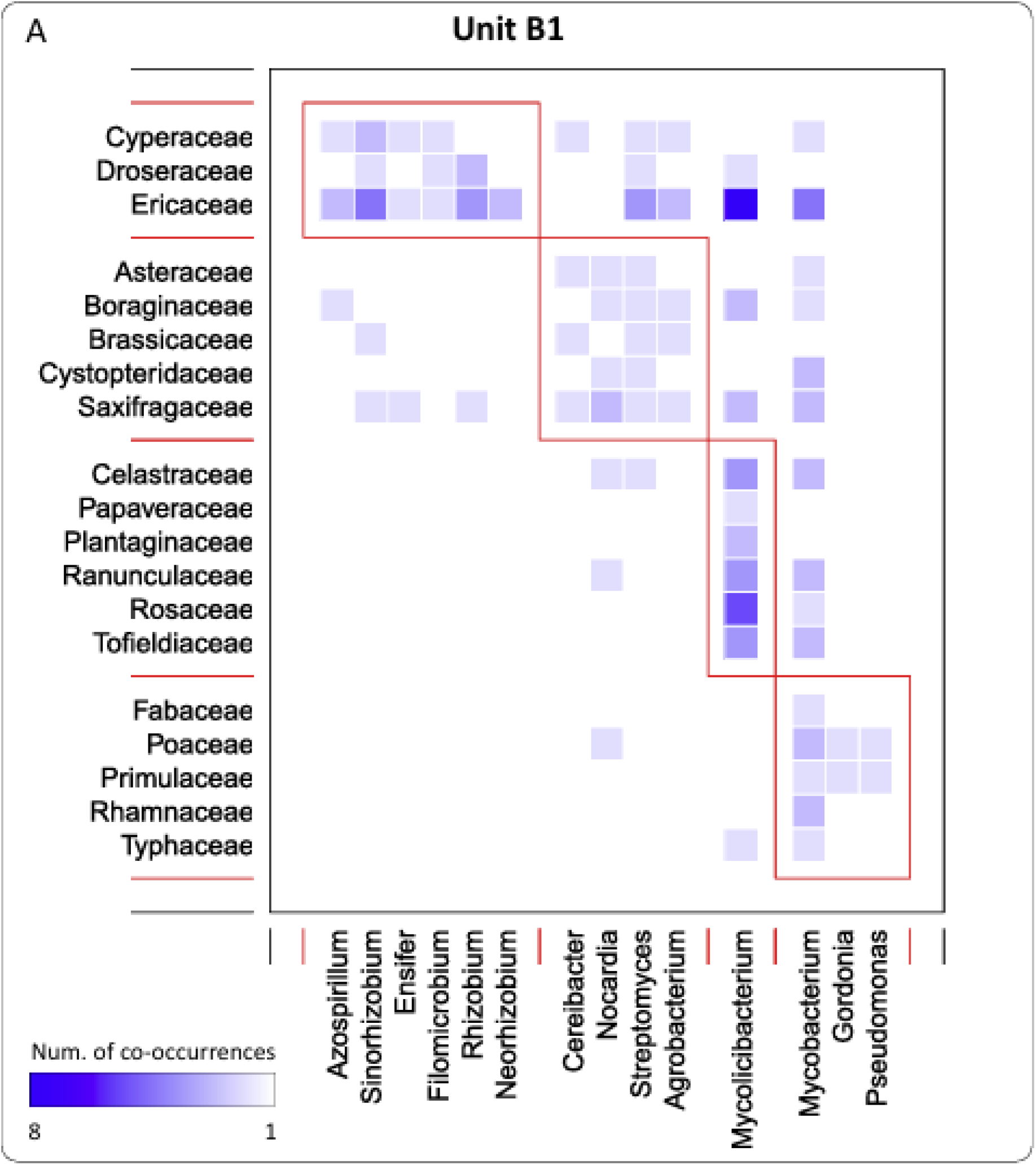
Plant–microbe co-occurrence modules identified in Unit B1. Heatmap showing the number of co-occurrences between plant families (rows) and microbial genera (columns). The intensity of the blue color indicates the number of co-occurrences. Red and black outlines denote the modules detected using the *computeModules* function (*bipartite* package, R environment), representing groups of plant and microbial taxa with higher within-module connectivity compared to the rest of the network.

**Figure 4.**
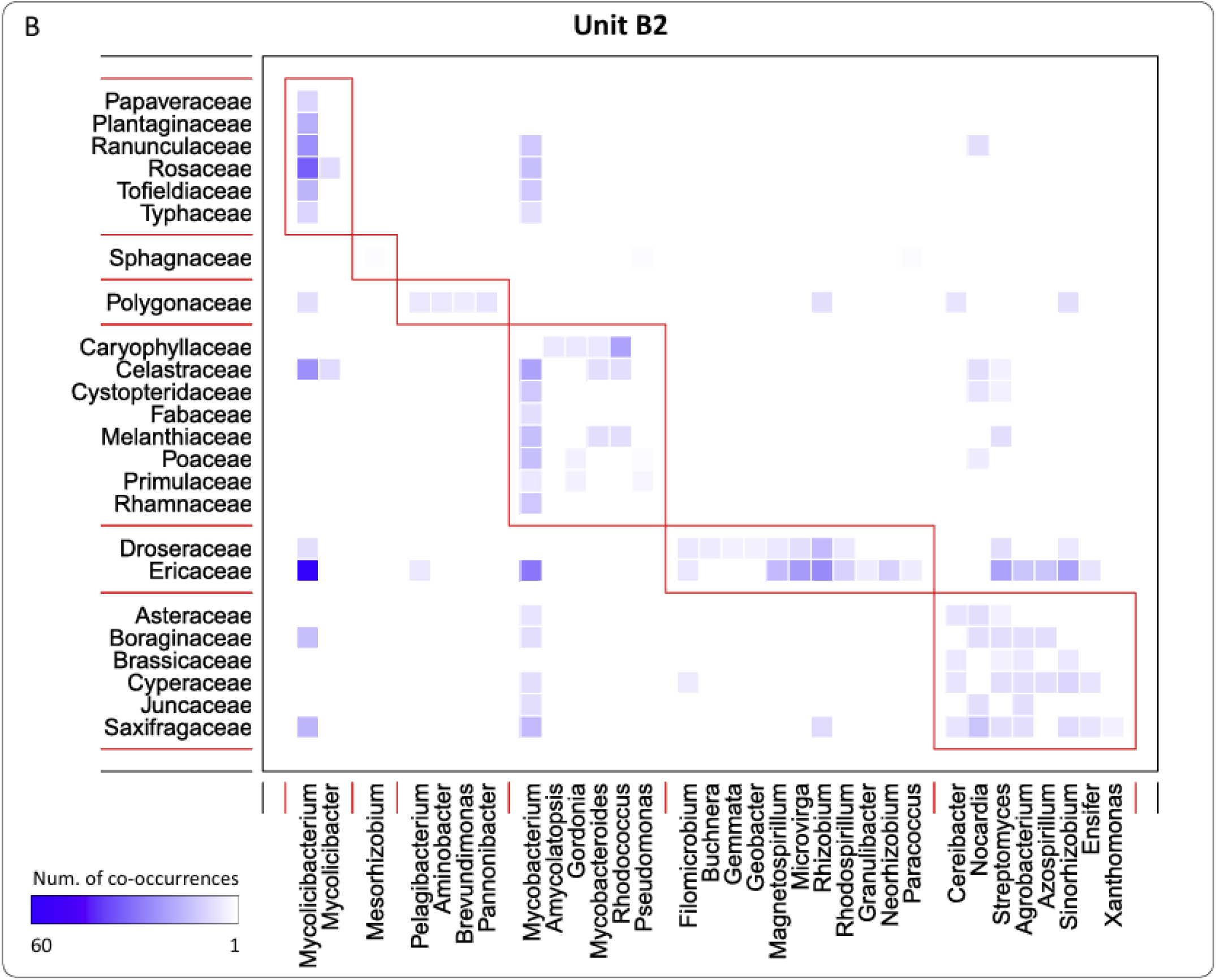
Plant–microbe co-occurrence modules identified in Unit B2. Heatmap showing the number of co-occurrences between plant families (rows) and microbial genera (columns). The intensity of the blue color indicates the number of co-occurrences. Red and black outlines denote the modules detected using the *computeModules* function (*bipartite* package, R environment), representing groups of plant and microbial taxa with higher within-module connectivity compared to the rest of the network.

**Figure 5.**
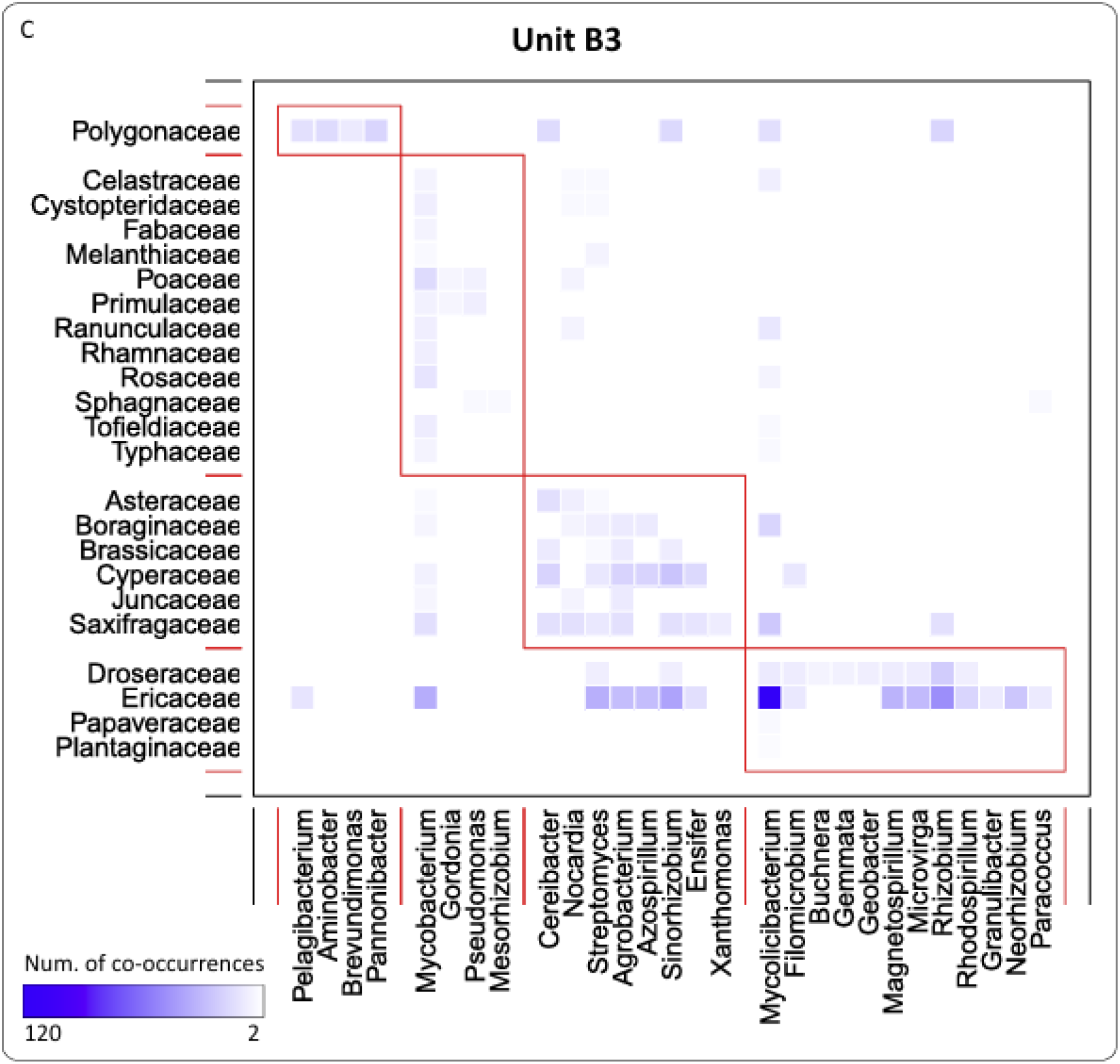
Plant–microbe co-occurrence modules identified in Unit B3. Heatmap showing the number of co-occurrences between plant families (rows) and microbial genera (columns). The intensity of the blue color indicates the number of co-occurrences. Red and black outlines denote the modules detected using the *computeModules* function (*bipartite* package, R environment), representing groups of plant and microbial taxa with higher within-module connectivity compared to the rest of the network.

By contrast, Unit B1 (Fig. 3) displayed more specialised patterns, characterised by weaker and less selective co-occurrences. Together, these observations suggest that environmental filtering and plant-specific niches may shape microbial community assembly differently across stratigraphic units, influencing both the strength and specificity of plant–microbe associations.

To further investigate the role of plant-growth-promoting bacteria (PGPB) in these ecosystems, we focused on a subset of microbial genera characterised by N-fixing ability and PGP traits, and typically found in the rhizosphere according to PlaBAse, to infer a rhizosphere-like component of the ancient microbial community. We constructed three bipartite networks, one for each stratigraphic unit, capturing only the co-occurrences between plants and these potential rhizospheric microorganisms. The resulting networks (Fig. 6) revealed a recurrent rhizosphere-like association core across stratigraphic units, suggesting that ancient sedaDNA can preserve ecological structure beyond taxonomic composition alone. Several plant families (such as Ericaceae, Cyperaceae, Saxifragaceae, Boraginaceae, and Brassicaceae) co-occurred with bacterial genera, including *Streptomyces, Agrobacterium, Azospirillum, Sinorhizobium*, and *Rhizobium*, across all units, forming a recurrent taxonomic core across the sampled stratigraphic units.

**Figure 6.**
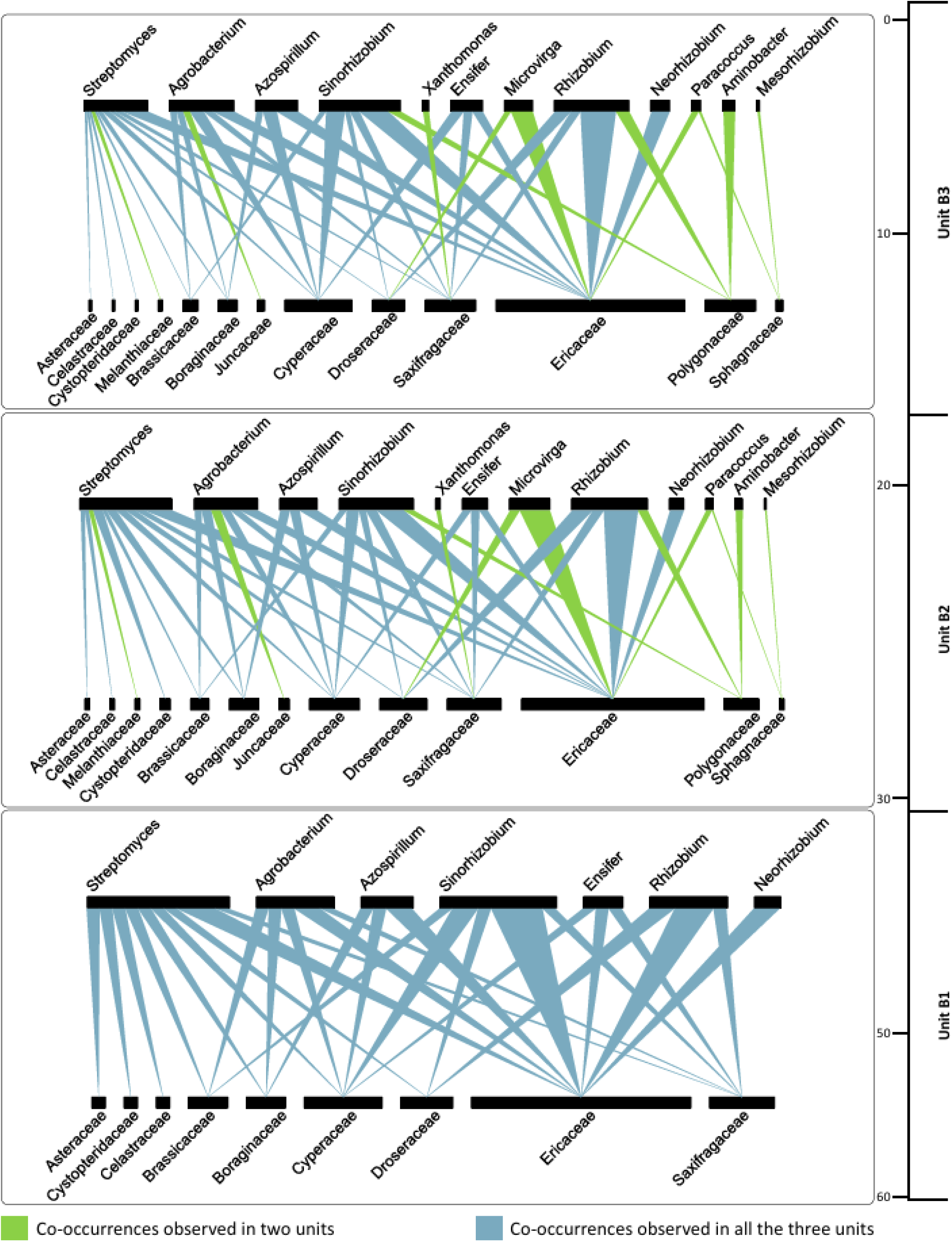
Bipartite networks showing the co-occurrences between the selected rhizosphere microorganisms and plants for each unit (Units B1, B2 and B3). Each panel shows a bipartite network connecting plant families (bottom nodes) and microbial genera (top nodes). The width of the links reflects the number of significant co-occurrences between a given plant–microbe pair

Interestingly, many of these associations reflect those found in modern plant–microbe interactions. For instance, several species of the Streptomyces genus can form well-documented relationships with grasses in the Boraginaceae and Asteraceae families and other monocots, promoting plant growth mainly through siderophore production and phosphate solubilization (Djemouai et al., 2022; Guo et al., 2021; Hamim et al., 2019). Likewise, *Rhizobium* and *Sinorhizobium* are classical N-fixing symbionts of legumes but are also known to engage non-nodulating relationships with non-leguminous hosts, including Ericaceae (Díez-Méndez & Menéndez, 2021; Hamim et al., 2019). These associations contribute to N cycling and improve stress tolerance in nutrient-poor soils (Santi et al., 2013).

Notably, Ericaceae, Droseraceae and Cyperaceae dominated the networks, emerging as key plant hosts, forming dense and repeated associations with N-fixing and siderophore-producing bacteria across all three units. These relationships reflect modern analogues, where sedges and ericaceous plants thriving in nutrient-poor environments actively recruit diazotrophic and PGP bacteria to support their growth (Carvalho et al., 2014; Lin et al., 2022; Mei et al., 2021; Rejmánková et al., 2018; Rilling et al., 2018). This pattern suggests that the observed microbial core recurs across the sampled stratigraphic units, revealing the potential of sedaDNA to detect repeated plant–microbe association signals in deep-time sediments.

The similarity of these associations to modern rhizosphere interactions suggests that some key plant–microbe relationships may have deep evolutionary roots, although direct continuity through time cannot be demonstrated from a single depositional system.

Taken together, these significant co-occurrence patterns indicate that ancient plant–microbe associations were detectable, consistent with H1, and support the novel potential of sedaDNA to infer conserved rhizosphere-like interaction networks in deep time.

Having detected recurrent, non-random plant–microbe associations, we next investigated whether the associated taxa harbour functional traits commonly involved in rhizosphere interactions and PGP potential. In line with H2, we identified across all three stratigraphic units a recurrent functional core composed of predicted genes associated with processes such as root colonisation, stress tolerance, and nutrient assimilation (Fig. 7). These traits are central to the establishment and maintenance of beneficial plant–microbe interactions and to the promotion of plant growth (Anas et al., 2025; Rosier et al., 2018). Remarkably, these same traits are still present in modern plant growth-promoting bacteria (Anas et al., 2025; Rosier et al., 2018), suggesting that the functional core is conserved not only spatially across all units but also temporally. This convergence with modern ecosystems highlights a key novel result of our study: sedaDNA can be used not only to reconstruct ancient community composition, but also to test whether taxonomic, network, and functional signals support rhizosphere-mediated plant in a 2-million-year-old ecosystem. These traits may therefore represent ancient adaptive solutions to plant–microbe symbiosis, preserved because of their ecological relevance and mutualistic benefits. Together, the persistence of both taxonomic and functional cores underscores the evolutionary importance of plant–microbe interactions and their crucial role in sustaining plant growth across timescales.

**Figure 7.**
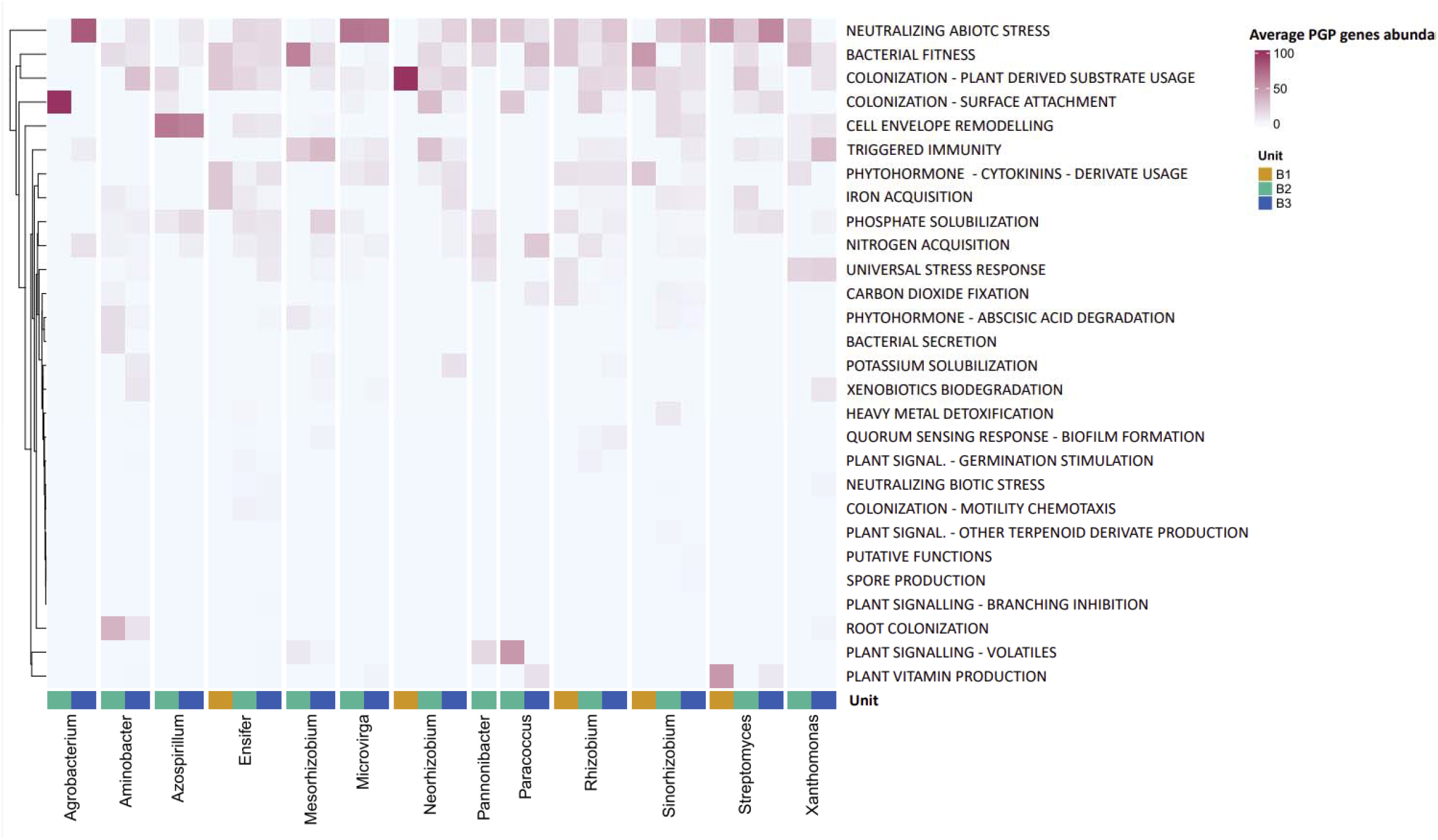
Functional potential of rhizospheric microbial communities based on PGPGFinder analysis. Heatmap showing the relative abundance (Z-score normalized counts) of plant growth-promoting genes (PGPGs) identified across microbial taxa within Units B1, B2, and B3 (Fig. 1B). Rows represent functional categories related to plant–microbe interactions (e.g., phytohormone production, nutrient acquisition, stress response, and colonization traits), and column represent microbial taxa.

However, several limitations should be considered when interpreting these results. First, plant– microbe associations were inferred from co-occurrence patterns in sedaDNA data, which cannot directly demonstrate ecological interactions or symbioses, as shared environmental filtering, sedimentary transport, and differential DNA preservation may also contribute to non-random co-distributions. Second, functional inferences are based on partially reconstructed microbial genomes and modern reference databases and therefore reflect potential rather than realised metabolic activity in the ancient ecosystem. Third, the analysis is limited to a single depositional system and time interval, restricting the ability to test temporal continuity or evolutionary conservation across independent climatic phases. Despite these constraints, the observed consistency of taxonomic, network, and functional patterns provides a conservative proof of principle that sedaDNA can capture structured, interaction-compatible ecological signals in deep time.

## Conclusions

Our findings support the interpretation that the boreal-like soils of the Kap København Formation functioned as complex ecosystems, in which microbial communities may have contributed to nutrient cycling and plant establishment in ways broadly comparable to modern soils. Our results show recurrent co-occurrence patterns between plant taxa and rhizosphere-associated microorganisms across the three stratigraphic units, supporting the presence of non-random, rhizosphere-compatible association signals and of a recurrent plant–microbe core across the sampled stratigraphic sequence (H1). The functional profiles of these bacteria, characterised by predicted plant-growth-promoting traits, further indicate a recurrent functional signal compatible with plant-associated microbial lifestyles, supporting our second hypothesis (H2). Most importantly, our analyses demonstrate that sedaDNA can address ecological questions beyond taxonomy, including whether ancient sediments preserve structured plant–microbe association signals. This integrated view of plant communities and their potentially associated bacteria offers new insight into how microbial networks may have contributed to ecosystem resilience and long-term environmental stability.

Together, these results highlight the potential of sedaDNA as a powerful complementary tool for investigating the ecological organisation of ancient ecosystems, while reinforcing the need for cautious interpretation when moving from co-occurrence to inferred interaction.

## Materials and methods

### Study site: Kap København Formation (Northern Greenland)

We analysed 2-million-year-old aDNA extracted from samples collected across four sites within the Kap København Formation in Northern Greenland (Fig. 1A) obtained from Kjær et al. (2022). All details regarding the extraction and sequencing procedures are provided therein. Briefly, the aDNA was was extracted from four sites of Member B of the Kap København Formation. Member B comprises 40–50 m of sandy (Units B1 and B3) and silty (Unit B2) deposits, including thin organic-rich beds with interglacial macrofossil assemblages deposited closer to the shore, within shallow marine to estuarine settings represented by upper and lower shoreface facies (Fig. 1B). A series of complementary studies has refined the depositional age of the Kap København Formation to an approximately 20,000-year interval around 2.4 Myr ago, during the Early Pleistocene. Nearly four decades of paleoenvironmental and climate research at the site indicate that, during this period, the now-desertic region was located within the boreal–Arctic ecotone, with summer and winter average minimum temperatures of approximately 10 °C and −17 °C, respectively, over 10 °C warmer than at present (Kjær et al., 2022). These relatively mild climatic conditions likely caused substantial ice-sheet ablation across Greenland, producing one of the last ice-free intervals in the past 2.4 million years (Funder et al., 2001). Fossil records indicate that during this interglacial period, the area supported a boreal forest mainly dominated by Ericaceae, Betulaceae, Salicaceae and Cyperaceae (Funder et al., 2001; Kjær et al., 2022). This is consistent with the vegetation reconstructed from plant and animal sedaDNA analyses reported by Kjær et al. (2022).

### Bioinformatic and network analysis

Raw sequencing reads were trimmed of Illumina adapters using BBDuk and overlapping paired-end reads were merged with BBMerge (Bushnell et al., 2017). The quality of pre-processed reads was assessed using FastQC (Simon Andrews et al., 2010). Taxonomic assignment was performed with KrakenUniq (Breitwieser et al., 2018), which employs a *k*-mer–based approach, using the NCBI GenBank (NT) database that includes microbial, vertebrate, non-vertebrate, and plant organisms. To minimise false-discovery rate, a taxon was considered detected only if it had at least 200 assigned reads and 1,000 assigned *k*-mers (Pochon et al., 2023). Additionally, all taxa detected in blanks and negative controls were excluded from the analysis.

To further validate taxonomic assignments, pre-processed reads were aligned via competitive mapping to an indexed NCBI GenBank (NT) database using Bowtie2 (Langmead & Salzberg, 2012), and the damage pattern for each detected organism was computed by MapDamage (v2.2.3) (Ginolhac et al., 2011). The read alignments were further filtered by evenness of coverage and presence of deamination, i.e. only microbial taxa that consistently exhibited convincing deamination signals (> 5% frequency of C/T polymorphisms at the ends of the aligned reads) and mean evenness of coverage profiles > 0.1% across all samples from a given site were considered authentic ancient organisms within that site. Taxonomic compositions were then analysed and visualised in R using the Phyloseq (McMurdie & Holmes, 2013) and ggplot2 (Wickham, 2016) packages.

For each alignment, we generated a consensus genome in FASTA format using BCFtools after restricting the analysis to reads identified as ancient. Variant discovery was performed with BCFtools mpileup (Danecek et al., 2021) against the sample⍰specific reference genome, and variants were called with *bcftools* call using the multiallelic caller (-m) while retaining variant sites only (-v). The resulting VCF file was compressed and indexed with *tabix*. To account for regions with insufficient support, depth as computed across the reference using *samtools* depth, and positions with coverage lower than 3x were converted into a BED mask. Consensus sequence were then generated with *bcftools consensus* by applying the called variants to the reference while masking low-coverage and uncovered positions. Functional annotation of plant growth⍰promoting genes was carried out with *PGPg_finder* (Pellegrinetti et al., 2024) using the *genome_wf* workflow. In this workflow, open reading frames are predicted with Prodigal and functionally annotated with DIAMOND (Buchfink et al., 2021) against the curated PLaBAse PGPT database, enabling the detection and categorisation of genes associated with plant growth⍰promoting traits, such as nutrient acquisition, stress tolerance, and signalling.

To evaluate the statistical significance of plant-bacteria co-occurrences, we applied Fisher’s exact test (Upton, 1992). Using the list of ancient plants reported by Kjær et al. together with the ancient microbial taxa identified in our analysis, we built a presence/absence matrix for each sample. In this matrix, plant taxa were represented by columns, microbial taxa by rows, and each cell contained either a 1 (taxon present) or a 0 (taxon absent). For each pairwise plant–microbe combination, we calculated a 2 × 2 contingency table: i) Cell a: number of samples in which both the plant and microorganism were present; ii) Cell b: number of samples in which the plant was present, but the microorganism was absent; iii) Cell c: number of samples in which the microorganism was present, but the plant was absent; iv) Cell d: number of samples in which neither the plant nor the microorganism was present. Fisher’s exact test was then applied to each contingency table to test the null hypothesis of independent distribution between plants and bacteria. To account for multiple comparisons, p-values were adjusted using the Benjamini–Hochberg false discovery rate (FDR) correction (Benjamini & Hochberg, 1995).

Associations with an FDR adjusted p-value < 0.05 were considered statistically significant. Co-occurrence can thereby be defined as the number of samples in which a given plant family and microbial taxon were jointly detected and passed FDR filtering and interpreted as statistical signals rather than direct evidence of physical interaction or symbiosis. For a better graphical representation and robustness, microbial taxa were grouped at the genus level, while plants were grouped at the family level. To specifically investigate potential rhizosphere-associated interactions, from the set of statistically significant co-occurrences identified through Fisher’s exact test, we selected only those involving microorganisms known to inhabit the rhizosphere, as indicated by the PLaBAse database (Patz et al., 2021). These filtered co-occurrences were then used to construct a bipartite network representing plant–microbe associations across all sampling sites. Network construction was performed using the R package *bipartite* (v2.21) (Dormann et al., 2008). To further characterize the potential plant–microbe interactions, microbial sequences from the statistically associated taxa were annotated using the *PGPg_finder* v1.1.0 workflow (Pellegrinetti et al., 2024) PLaBAse database, which provides unique information on genes implicated in plant–microbe interactions, including those related to N fixation and other functional traits associated with plant growth-promoting rhizobacteria (PGPR) potential.

## Data availability

The data supporting the findings of this study have been deposited in Zenodo under accession DOI https://doi.org/10.5281/zenodo.20928857.

## Acknowledgements

This work was supported by a grant from the Free University of Bozen/Bolzano through the Start-up Fund (AG2803 - EARTHCODE), Infra 2025 (PH2001 - BIOAGRI-HPC) and the European Union – NextGenerationEU. The authors thank Eske Willerslev, Antonio Fernandez-Guerra and Kurt Kjær for their valuable support.

NO is financially supported by the TARGETWISE project (Functional Omics Analysis of Metabolic Diseases to Advance Drug Discovery Research Excellence in LIOS, project No. 1.1.1.5/2/24/A/003, Latvian Institute of Organic Synthesis, 2024–2029; https://targetwise.osi.lv/).

